# GPU-accelerated mesh-based Monte Carlo photon transport simulations

**DOI:** 10.1101/815977

**Authors:** Qianqian Fang, Shijie Yan

## Abstract

The mesh-based Monte Carlo (MMC) algorithm is increasingly used as the gold-standard for developing new biophotonics modeling techniques in 3-D complex tissues, including both diffusion-based and various Monte Carlo (MC) based methods. Compared to multi-layered and voxel-based MCs, MMC can utilize tetrahedral meshes to gain improved anatomical accuracy, but also results in higher computational and memory demands. Previous attempts of accelerating MMC using graphics processing units (GPUs) have yielded limited performance improvement and are not publicly available. Here we report a highly efficient MMC – MMCL – using the OpenCL heterogeneous computing framework, and demonstrate a speedup ratio up to 420× compared to state-of-the-art single-threaded CPU simulations. The MMCL simulator supports almost all advanced features found in our widely disseminated MMC software, such as support for a dozen of complex source forms, wide-field detectors, boundary reflection, photon replay and storing a rich set of detected photon information. Furthermore, this tool supports a wide range of GPUs/CPUs across vendors and is freely available with full source codes and benchmark suites at http://mcx.space/#mmc.

## 1 Introduction

Modeling photon-tissue interactions accurately and efficiently is essential for an array of emerging biophotonics applications, such as diffuse optical tomography (DOT), fluorescence molecular tomography (FMT) and functional near-infrared spectroscopy (fNIRS). Due to the complex nature of photon-tissue interactions, significant effort has been dedicated towards developing computationally efficient methods that not only properly consider the underlying physical processes, such as light scattering, absorption and emission, but also accurately model the complex-shaped anatomical boundaries that delineate tissues.

Over the past decade, Monte Carlo (MC) based modeling has seen increasing use, thanks to two recent breakthroughs in MC algorithm development. The first breakthrough takes advantage of the parallel computing capability of modern graphics processing units (GPUs) and dramatically shortens the computational time by 2 to 3 orders of magnitude.^1, 2^ The second breakthrough allows MC to model complex tissue boundaries using increasingly sophisticated representations, such as 3-D voxels,^1, 3^ non-uniform grids, and triangular^4^ and tetrahedral meshes.^5, 6^ Combined with the generality and intuitive domain settings, these new advances has made MC not only a choice for gold-standard solutions, but also a powerful research tool increasingly involved in routine optical data processing and instrument parameter optimizations.

Nearly a decade ago, our group proposed one of the first mesh-based MC (MMC) methods.^6^ Compared with the traditional voxel-based MC, MMC reports better accuracy because meshes are more anatomically accurate in modeling arbitrarily-shaped 3-D tissues, which are often delineated by curved boundaries. Since then, a number of works have been published to further extend MMC for faster and more accurate simulations. In several works,^7, 8^ the ray-tracing calculation and random number generation were vectorized and ported to single-instruction multiple-data (SIMD). This data level parallelism significantly improves the simulation speed on modern CPUs. Yao *et al.*^9^ reported a generalized MMC to efficiently support wide-field illumination and camera-based detection. In Yan *et al.*,^10^ a dual-grid MMC, where a coarsely tessellated mesh and a fine grid are used for ray-tracing and output storage, respectively, managed to enhance both performance and solution accuracy. Leino *et al.*^11^ developed an open-source MMC software which incorporates MATLAB interface for improved usability.

Despite the steadily growing user community, a highly efficient GPU-accelerated MMC implementation remains missing. While there has been a number of attempts to accelerate MMC using GPU computing, only limited success has been reported. For example, Powell *et al.*^12^ reported a CUDA-based GPU-MMC for acoustic-optics (AO) modeling. The authors reported that, in the context of different application focuses, an ~8x speedup was achieved when comparing their GPU algorith with our single-threaded MMC simulation. In another work,^13^ Zoller *et al.* reported a GPU-based MMC (MCtet) to model anisotropic light propagation in aligned structures. Unfortunately, the software codes in both works are not publicly available to allow further testing or comparison. While these results are encouraging, compared to highly accelerated voxel-based MC,^1, 14^ the lower magnitude in speedup presents an opportunity for further improvement.

The difficulties of accelerating MMC in modern GPU processors are associated with both the computation and memory characteristics of MMC photon modeling. Compared to voxel- and layer-based MC algorithms, MMC requires more geometric data (mesh nodes, elements, and surfaces) to advance a photon per propagation step. Because GPUs typically have scarce high-speed shared memory and constant memory,^15, 16^ the memory latency becomes a major barrier towards allowing MMC to benefit from the parallel hardware. Also, the ray-tracing calculations in MMC involve more complex tests to calculate ray-triangle intersections within the enclosing tetrahedra. This imposes higher computational demands compared to MC models that only handle simple geometries.

Here, we report a highly accelerated MMC implementation, MMCL, developed using the Open Computing Language (OpenCL) framework. MMCL supports almost all advanced simulation features seen in our open-source CPU MMC, such as the support of wide-field complex sources and detectors,^9^ photon replay,^17^ dual-grid simulations^10^ and storage capabilities for rich sets of detected photon data. Thanks to the excellent portability of OpenCL, the MMCL is capable of running on a wide range of commodity GPUs. From our tests, MMCL has shown about 2 orders of magnitude speedup compared to our highly optimized single-thread CPU implementation. Although we recognize the 2- to 5-fold speed disadvantage for OpenCL compared to CUDA on NVIDIA hardware, as shown in our previous study,^14^ in this work, our decision of prioritizing the development of OpenCL MMC is largely motivated by 1) the open-source ecosystem of OpenCL allowing the developed software to be rapidly and widely disseminated through public software repositories, and 2) the upcoming high-performance GPUs from Intel and AMD being expected to attract more development attention to OpenCL libraries and drivers, likely resulting in boosts in performance.

In the following sections, we will first discuss the key algorithm steps and optimizations that enable high-throughput MC simulations, and then report our validation and speed benchmarks using simulation domains covering a wide range of complexities and optical properties. Finally, we discuss our plans to further develop this technique.

## 2 Methods

### 2.1 GPU-accelerated photon propagation in tetrahedral meshes

As we discussed previously,^7^ at the core of MC light transport modeling is a ray-tracing algorithm that propagates photons through complex media. In our publicly available MMC software,^6^ we have implemented 4 different ray-tracers to advance photons from one tetrahedron to the next. These ray-tracers are based on 1) the Plücker coordinates,^6^ 2) a fast SIMD based ray-tracer,^7^ 3) a Badouel ray-tracing algorithm,^5, 18^ and 4) an SIMD-based branchless-Badouel ray-tracer.^19^ As we demonstrated before,^7^ the branchless-Badouel ray-tracing algorithm reported the best performance among the above methods; it also requires the least amount of memory and computing resources. For these reasons, we specifically choose the branchless-Badouel ray-tracer in this work.

In our CPU based MMC software, we explored SIMD parallelism using SIMD Extensions 4 (SSE4).^7^ In this work, we ported our manually tuned SSE4 computation to OpenCL, resulting in both improved code-readability and efficiency. The ray-tracing calculation in our GPU MMC algorithm can be represented by the below 5-step 4-component-vector operations:

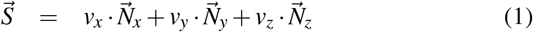

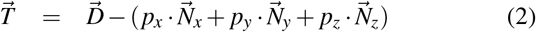

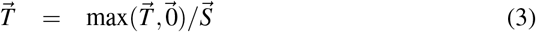

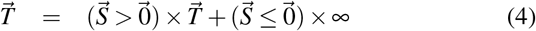

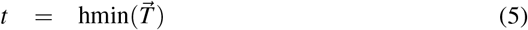

where ×, /, `max` are element-wise multiplication, division and maximum value, respectively; 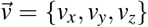 and 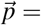 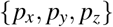 are the current photon direction and position, respectively; 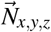 denotes the *x*/*y*/*z* components (4 elements in each vector) of the surface normal vectors 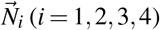 at the 4 facets of the current tetrahedron; 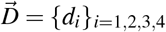 is computed by 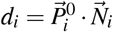, where the 3-D position 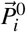 can be any node of the *i*-th face as the dot product with 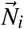 is a constant per face;^18^ 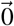 is an all-zero vector; and `hmin` performs the “horizontal” minimum to extract the lowest value from the 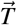 vector. Both 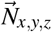 and 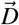 are pre-computed. Vectors 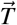 and 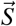 in Eqs. 1 to 4 are intermediate variables, and 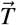 in Eq. 5 denotes the distances from 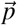 to the intersection points of the 4 facets of the current tetrahedron (intersections in the 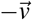 direction are ignored). The index in 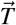 where the distance has the lowest value, i.e. *t* in Eq. 5, indicates the triangle that the ray intersects first. Notice that above calculations consist of only short-vector (3 or 4 elements) operations with no branching. Such calculations can be efficiently optimized on the modern GPUs or CPUs, resulting in high computational throughput.

The above vectored formulation is also illustrated in Fig.1. The “photon ray” with origin 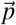 and direction 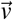 intersects with the 4 facets at 4 distances, *t*_*i*_: a positive distance indicates intersection in the forward direction and a negative distance indicates an intersection in the 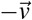 direction. The task of finding the intersection point becomes finding the minimum positive distance in 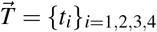. This is efficiently achieved by first replacing all negative values in 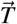 by +∞ in Eq. 4 and then taking a horizontal minimum in Eq. 5.

**Fig. 1.**
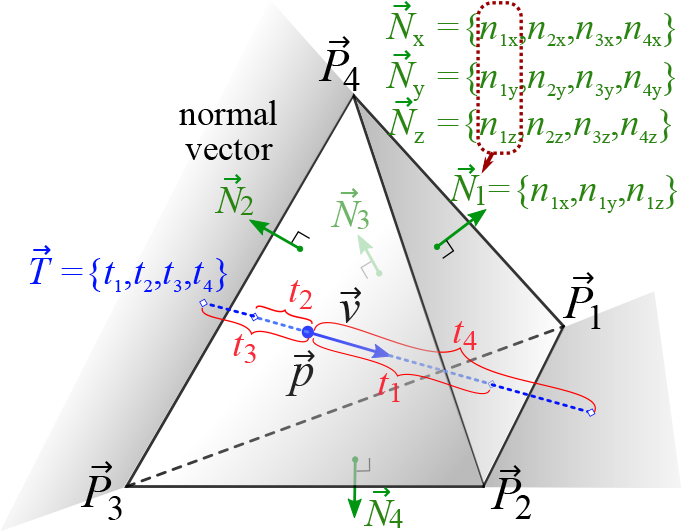
Illustration of a ray-tetrahedron intersection testing using the branchless Badouel algorithm. 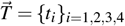 records the signed distances from the current position 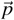 to the 4 facets of the tetrahedron along the current direction 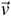. A negative distance means intersecting in the 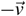 direction (such as *t*_2_ and *t*_3_).

The above MMCL algorithm readily supports a variety of wide-field sources and detectors via our mesh retesselation approach.^9^ By storing various photon-packet related parameters, such as partial-pathlength, scattering event count, momentum transfer etc, MMCL is also capable of exporting a rich set of detected photon information, similar to its CPU counterpart. Our previously developed “photon replay” approach^17^ for constructing the Jacobian matrix is also implemented in this OpenCL code.

### 2.2 Dual-grid MMC GPU optimization

Recently, we reported a significantly improved MMC algorithm, the dual-grid MMC or DMMC,^10^ combining the strengths of both voxel- and mesh-based simulations. It reduces the ray-tracing overhead significantly by utilizing a coarse tessellation of the surface nodes without losing anatomical accuracy while a dense voxelated storage grid provides higher spatial resolution and lower discretization error. In this work, we have successfully ported the DMMC algorithm from SIMD to an OpenCL-based computing model. The outputs are written in a voxelated grid in the GPU/CPU, similar to our voxel-based MC methods.^14^ To reduce global memory operations, we only write to memory when a photon packet attempts to move out of a voxel in the output grid.

### 2.3 GPU memory characterization and optimization

Compared to voxel-based Monte Carlo algorithms, MMC has different memory characteristics. These can greatly impact the computational efficiency on GPUs because high speed memory is limited on GPUs. To advance a photon packet by one step in a tetrahedral mesh, MMC requires reading more geometric data, including node coordinates, node indices of the current element, normal vectors of the tetrahedron facets etc. Although most of these data can be pre-computed on the host (CPU) and copied to the GPU, such data are too big to be stored into the high-speed shared or constant memory. Therefore, one has to store the bulk of the mesh data in the “global memory” which is known to have high latency (roughly 100× slower than the shared memory).^16^ It is important to minimize such latency for an efficient GPU MMC implementation.

The key to overcoming this memory latency issue on the GPU is to launch a large number of threads and ensure that the GPU streaming multi-processors (SM) have abundant active thread-blocks (divided into “warps” in the NVIDIA literature and “wavefronts” in OpenCL literature). Therefore, while some of the threads wait for data from the global memory, the GPU scheduler can switch on other threads and keep the SMs busy. When there are sufficient wavefronts in the waiting queue, the global memory latency can be effectively hidden. Typically, a minimum of 10-20 wavefronts per SM is required^15, 16^ to effectively hide the memory latency on the NVIDIA and AMD GPUs. However, the maximum wave-fronts per SM is strongly dependent on how the kernel is programmed, especially in regards to the number of registers and size of shared memory as SMs have only limited resources and must share them among all active wavefronts. As a result, the key to accelerate the GPU based MMC is to 1) launch a large number of threads that produce sufficiently large waiting queues per SM, and 2) minimize the register/shared memory needs per thread so that many wave-fronts can simultaneously run on the SM. Following these insights, we are able to dramatically improve the simulation speed on tested GPU devices, independent of its vendors.

### 2.4 Simulations on heterogeneous computing platforms

As discussed in our previous investigation,^14^ the high scalability of OpenCL permits simultaneous use of multiple computing devices. This includes both simultaneous use of different generations of GPU architecture and mix between GPUs and CPUs. In such a heterogeneous computing environment, it is crucial to ensure that the simulation algorithm has a flexible workload distribution strategy to assign appropriate workloads to each GPU device according to their capabilities.

Several device-level load-balancing strategies have been implemented in this work. A manual workload distribution can be specified by a user. The manual workload partition can be guided using relative speeds obtained from a small workload running on each device. In addition, we also provide a heuristic method to automatically determine an effective workload partition according to the persistent thread (PT)^20^ count on each invoked device. The PT count is determined by the number of threads that can “fully occupy” all SMs on the device. For example, for an Intel GPU, the PT count is computed by block size × 7 × #EU (execution unit, 1 EU can run 7 threads^21^); for AMD GPUs, we estimate PT count by block size × 40 × #CU (compute unit, 1 CU can run 40 active wavefronts^16^).

## 3 Results

In this section, we first validate our massively parallel MMCL algorithm using standard benchmarks and then systematically characterize and compare speedup ratios over a range of CPU/GPU devices using single- and multi-threaded MMC on CPUs as references. The simulation speeds are reported for both conventional single-grid and dual-grid MMC (DMMC). For all simulations, 10^8^ photons are simulated with atomic operations^1^ enabled. All benchmarks were performed on Ubuntu Linux 16.04 with the latest stable GPU drivers. All simulation input data and scripts are provided in our open-source code repository on Github (http://github.com/fangq/mmc) for reproducibility and future comparisons.

In the first set of benchmarks, we focus on validating the accuracy of MMCL in two simple geometries: **B1** (cube60) – a cubic homogeneous domain (see Fig. 2 in Ref.^6^) and **B2**(sphshells) – a heterogeneous domain made of multi-layered spherical shells (Fig. 1c in Ref.^10^).

**Fig. 2.**
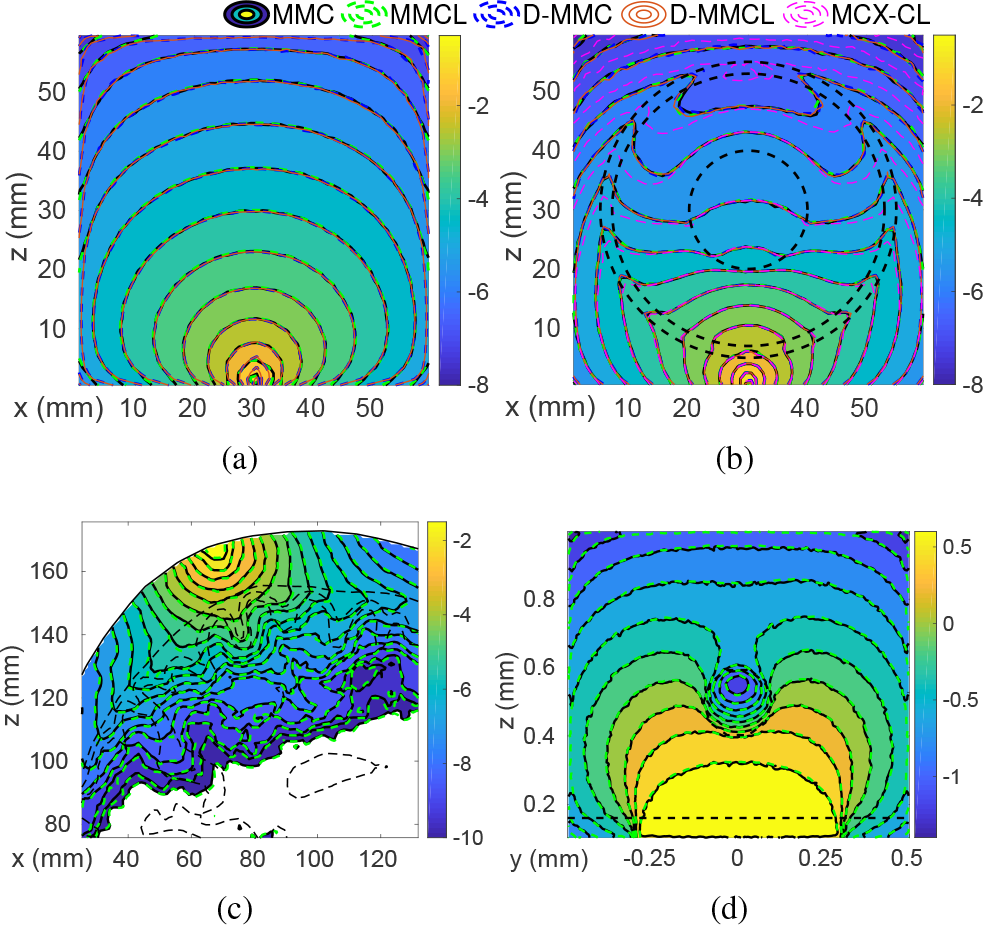
Fluence (mm^−2^, in log 10-scale) contour plots of MMC and MMCL in various benchmarks: (a) B1/B1D, (b) B2/B2D, (c) B3 and (d) B4. In (b), we also include a voxel-based MC (MCX-CL) output for comparison. Black-dashed-lines mark tissue boundaries.

Briefly, the **B1** benchmark contains a 60 × 60 × 60 mm^3^ homogeneous domain with an absorption coefficient *μ*_*a*_ = 0.005 mm^−1^, scattering coefficient *μ*_*s*_ = 1 mm^−1^, anisotropy *g* = 0.01, and refractive index *n* = 1.0. The medium outside of the cubic domain is considered as air. In the **B2** benchmark, the optical properties and dimensions of each layer are described previously.^10^ In both cases, a pencil beam source injects photons at [30, 30, 0] mm, with the initial direction pointing along the +*z*-axis. Boundary reflection is considered in **B2** but not in **B1**.

Both MMC and MMCL share the same tetrahedral mesh for each simulation. For **B1** and **B2**, two mesh densities were created to separately compare MMC and MMCL in the single-grid and dual-grid^10^ (denoted as **B1D** and **B2D**) simulation methods. The node and element numbers for the generated meshes along with the pre-accelerated MMC simulation speeds are summarized in Table 1. We want to mention that MMC simulation speed is correlated more strongly with the mesh density (normalized by the local scattering coefficient) within the high-fluence regions than the total mesh element/node sizes. For a fixed domain, however, increased mesh density results in higher element/node numbers.

**Tab. 1.** Summary of the meshes and baseline simulation speeds (in photon/ms, the higher the faster) for the selected benchmarks. The baseline speeds were measured using a single-thread (MMC-1) or 8-thread (MMC-8) SSE4-enabled MMC on an i7-7700K CPU.

| Benchmark | B1<br>(cube60) | B1D<br>(d-cube60) | B2<br>(sphshells) | B2D<br>(d-sphshells) | B3<br>(colin27) | B4<br>(skin-vessel) |
| --- | --- | --- | --- | --- | --- | --- |
| Node# | 29,791 | 8 | 604,297 | 3,723 | 70,226 | 76,450 |
| Elem.# | 135,000 | 6 | 3,733,387 | 21,256 | 423,375 | 483,128 |
| MMC-1 | 67.25 | 100.06 | 5.73 | 10.39 | 12.34 | 36.72 |
| MMC-8 | 351.26 | 568.06 | 26.43 | 57.49 | 67.43 | 150.42 |

In Fig. 2(a), we show the cross-sectional contour plots of the fluence distributions (mm^−2^) in log 10-scale using the fine-mesh model along the plane *y* = 30.5 mm. We also overlap the result from the DMMC output with those from MMCL to show the agreements between different MC methods.

In the 2^nd^ set of tests, we expand our comparisons to more challenging cases involving realistic complex domains. Two simulations are compared: **B3** (colin27) – simulations on a complex brain atlas – Colin27 (Fig. 4 in^6^), and **B4** (skin-vessel) – the skin-vessel benchmark.^22^ Briefly, the **B3** benchmark contains a 4-layer brain mesh model derived from an atlas;^6^ **B4** contains a 3-layer skin-model with an embedded vessel. The generated meshes for the two complex examples are also summarized in Table 1. The optical properties for **B3** are described in Ref.,^6^ and those for **B4** are described in Ref.^22^

In Fig. 2(c-d), we show the cross-sectional contour plots from the two complex cases. For **B3**, we show the sagittal slice along the source plane; for the **B4** benchmark, the cross-section is perpendicular to the *y*-axis aligned vessel. The curved tissue boundaries are shown as black-dashed lines.

Next, our focus is to benchmark the simulation speeds across a wide range of CPU/GPU processors using MMCL, and compare those with the baseline, i.e. single-threaded (“MMC-1” in Table 1) or multi-threaded (“MMC-8” in Table 1) SSE4-enabled MMC on the CPU. The branch-less Badouel ray-tracing algorithm^7, 19^ and a parallel `xorshift128+`^23^ random number generator are used across all MMC and MMCL simulations.

In Fig. 3(a), we plot the speedup ratios over single-threaded MMC (MMC-1) across a list of benchmarks and processors. In Fig. 3(b), we also included a comparison to MCX-CL^14^ – a voxel-based MC software using OpenCL. The voxelated simulation domain in MCX-CL matches the DMMC output grid of the corresponding MMCL simulations. The simulation speed numbers (in photon/ms) corresponding to Fig. 3(a) are also summarized in Table 2.

**Fig. 3.**
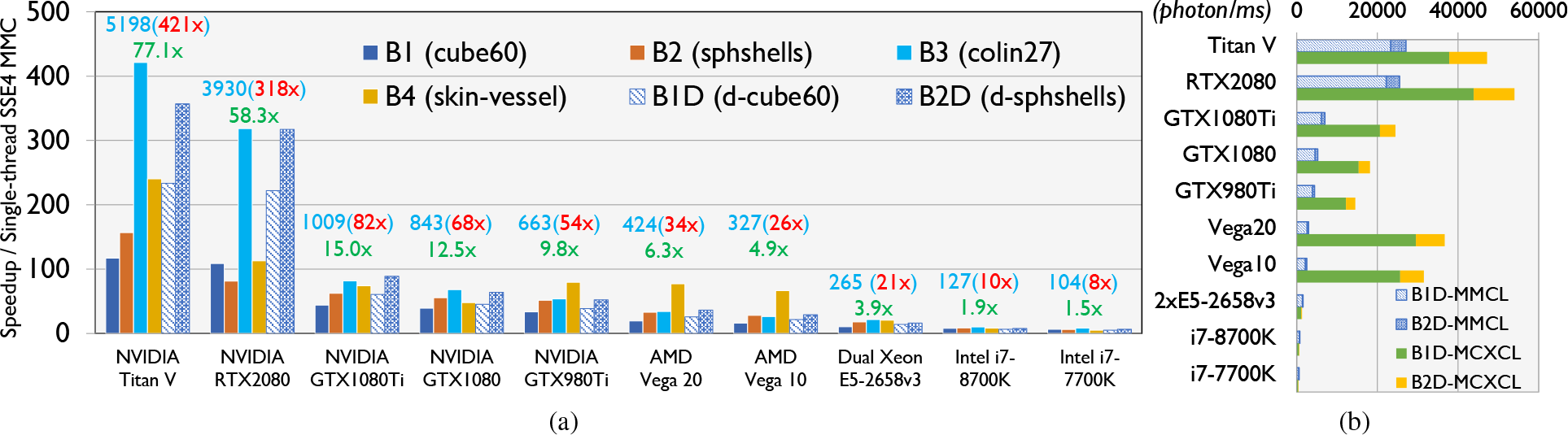
Speeds of MMCL in 6 benchmarks (B1-B4: single-grid; B1D/B2D: dual-grid): (a) speedup ratios over a single-threaded (on i7-7700K) SSE4 MMC, and (b) speeds in dual-grid simulations compared to MCX-CL. In (a), we also report the speed (photon/ms, light-blue) and speedups over the single-(red) and multi-threaded (green) MMC in the labels for Benchmark-B3 (Colin27).

**Tab. 2.** Simulation speed (in photon/ms, the higher the faster) of MMCL in 6 benchmarks (B1-B4: single grid; B1D and B2D: dual-grid); we also report the voxel-based MCX-CL speed in benchmarks B1D and B2D. The master script to reproduce the above results can be found in the “mmc/examples/mmclbench/” folder of our software.

| Device | B1 | B2 | B3 | B4 | B1D | B2D | B1D (MCXCL) | B2D (MCXCL) |
| --- | --- | --- | --- | --- | --- | --- | --- | --- |
| NVIDIA Titan V | 7874.02 | 899.25 | 5198.05 | 8829.24 | 23359.03 | 3709.20 | 37835.79 | 9353.66 |
| NVIDIA RTX 2080 | 7319.04 | 465.62 | 3930.20 | 4147.83 | 22202.49 | 3295.87 | 43917.44 | 10095.91 |
| NVIDIA GTX 1080Ti | 2959.81 | 357.89 | 1008.48 | 2721.38 | 6079.03 | 924.16 | 20648.36 | 3826.14 |
| NVIDIA GTX 1080 | 2642.92 | 318.22 | 843.28 | 1759.82 | 4547.73 | 665.72 | 15351.55 | 2796.73 |
| NVIDIA GTX 980Ti | 2254.44 | 295.36 | 663.21 | 2925.43 | 3872.37 | 542.73 | 12211.50 | 2319.06 |
| AMD Vega 20 | 1315.62 | 189.17 | 424.07 | 2839.54 | 2579.51 | 378.00 | 29577.05 | 7105.30 |
| AMD Vega 10 | 1086.45 | 161.42 | 326.89 | 2444.87 | 2170.52 | 302.46 | 25680.53 | 5865.45 |
| Dual Xeon E5-2658v3 | 708.19 | 103.86 | 265.00 | 756.74 | 1405.90 | 167.69 | 1127.86 | 181.83 |
| Intel i7-8700K | 538.80 | 49.29 | 126.80 | 313.77 | 670.12 | 80.92 | 597.07 | 89.02 |
| Intel i7-7700K | 434.86 | 35.41 | 103.64 | 188.83 | 522.77 | 67.28 | 397.16 | 58.96 |

In addition, we also test MMCL using multiple GPU devices simultaneously. Two AMD GPUs of different computing capabilities – Vega10 (Vega64) and Vega20 (Vega II) – are used. When running the **B1D** test on both GPUs, we distributed the workload based on the ratio between their single-GPU speeds. When using both Vega GPUs, we have obtained a speed of 4,583 photon/ms in the **B1D** benchmark; the speeds using Vega10 and Vega20 alone are 2,171 and 2,580 (in photon/ms), respectively.

### 3.1 Discussions and Conclusion

As expected, the contour plots generated from MMC and MMCL are nearly indistinguishable from each other in all four tested cases in Fig. 2. Similarly, in Figs. 2(a-b), the dual-grid MMC (DMMC) and MMCL (D-MMCL) outputs also excellently match the single-grid outputs. In Fig. 2(b), The previously observed fluence differences inside the spherical shell between mesh-based simulations and voxel-based MC simulations (MCX-CL)^10^ are also reproduced in MMCL outputs, emphasizing the importance of using mesh models when simulating domains with curved boundaries and high-contrast heterogeneities.

Many observations can be made from the speed results reported in Fig. 3(a). First, the high scalability of our GPU accelerated MMC algorithm is indicated by the increasing speedups obtained from more recent and capable GPUs. The simulation speed achieved on the NVIDIA Titan V GPU is 117× to 421× higher than that of the CPU-based MMC on a single thread; the highest acceleration on AMD GPUs is achieved on the Vega20 GPU, reporting a 20× to 77× speedup. Moreover, the high portability of the algorithm is evident by the wide range of NVIDIA/AMD/Intel CPUs and GPUs tested, owing to OpenCL’s wide support. There is a steady and significant increase in computing speed between different generations of GPUs made by these vendors, similar to our previous findings in the voxel-based MC algorithm.^14^

Before comparing the speed differences in mesh and voxel-based simulations, as shown in Fig. 3(b), one must be aware that these two algorithms result in differing levels of accuracy, as suggested in Fig. 2(b), even when using the same grid space for output. Nevertheless, several interesting findings can be drawn from this figure. In all NVIDIA GPUs, MMCL is about 30% to 50% of the speed compared to voxel-based MCX-CL.^14^ On AMD GPUs, MMCL is only ~8% of MCX-CL’s speed. However, Intel CPUs show an opposite result – MMCL is about 12% to 31% faster than MCX-CL in **B1D**. The sub-optimal speed on AMD GPUs is also noticeable in Fig. 3(a), where the speedup from Vega20 is only a fraction of the speedup from NVIDIA RTX2080, despite the former having 30% higher theoretical throughput.

To understand this issue further, we performed a profiling analysis and discovered that the sub-optimal speed on AMD GPUs results from extensive register allocation by the compiler – the AMD compiler produces ~200 vector registers compared to only 69 by the NVIDIA compiler. The high register count limits the total active blocks to only 4 on Vega20 compared to 28 on NVIDIA GPUs, making it extremely difficult for the GPU to “hide” memory latency.^16^ We are currently collaborating with AMD to investigate this issue.

Moreover, based on our previous observations in voxel-based MC using OpenCL and CUDA on NVIDIA devices,^14^ we noticed that CUDA-based MC simulation is about 2-to-5-fold faster than the OpenCL implementation due to driver support differences. As a result, we anticipate that if we further port MMCL to the CUDA programming language, one may achieve further speed improvement, with the resulting software limited to NVIDIA GPUs only.

In summary, we report a massively-parallel mesh-based Monte Carlo algorithm that offers a combination of both high speed and accuracy. The OpenCL implementation allows one to run high-throughput MC photon simulations on a wide range of CPUs and GPUs, showing excellent scalability to accommodate increasingly more powerful GPU architectures. We describe the insights that we have learned regarding GPU memory utilization and vendor differences. In addition, we provide speed benchmarks ranging from simple homogeneous domains to highly sophisticated real-world models. We report the speed comparisons between CPUs and GPUs made by AMD, NVIDIA and Intel, and show excellent portability between different devices and architectures. Our accelerated open-source software, including MATLAB/Octave support, is freely available at http://mcx.space/#mmc.

## Acknowledgments

This research is supported by the National Institutes of Health (NIH) grants R01-GM114365, R01-CA204443 and R01-EB026998. We would like to thank Dr. Samuel Powell for his helpful comments on comparisons with literature results.

**Qianqian Fang**, PhD, is currently an assistant professor in the Bioengineering Department, Northeastern University, Boston, USA. He received his PhD degree from Dartmouth College in 2005. He then joined Massachusetts General Hospital and became an Instructor of Radiology in 2009 and Assistant Professor of Radiology in 2012, before he joined Northeastern University in 2015. His research interests include translational medical imaging devices, multi-modal imaging, image reconstruction algorithms, and high performance computing tools to facilitate the development of next-generation imaging platforms.

**Shijie Yan** is a doctoral candidate at Northeastern University. He received his BE degree from Southeast University, China, in 2013 and MS from Northeastern University in 2017. His research interests include Monte Carlo photon transport simulation algorithms, parallel computing, GPU programming and optimization.

## References

1. Q. Fang and D. A. Boas, “Monte Carlo simulation of photon migration in 3D turbid media accelerated by graphics processing units,” Opt. Express 17(22), 20178–20190 (2009).

2. E. Alerstam, W. C. Y. Lo, T. D. Han, et al., “Next-generation acceleration and code optimization for light transport in turbid media using GPUs,” Biomed. Opt. Express 1(2), 658–675 (2010).

3. D. Boas, J. Culver, J. Stott, et al., “Three dimensional Monte Carlo code for photon migration through complex heterogeneous media including the adult human head,” Optics Express 10(3), 159–170 (2002).

4. N. Ren, J. Liang, X. Qu, et al., “GPU-based Monte Carlo simulation for light propagation in complex heterogeneous tissues,” Opt. Express 18(7), 6811–6823 (2010).

5. H. Shen and G. Wang, “A tetrahedron-based inhomogeneous Monte Carlo optical simulator,” Phys. Med. Biol. 55(4), 947–962 (2010).

6. Q. Fang, “Mesh-based Monte Carlo method using fast raytracing in Plücker coordinates,” Biomed. Opt. Express 1(1), 165–175 (2010).

7. Q. Fang and D. Kaeli, “Accelerating mesh-based Monte Carlo method on modern CPU architectures,” Biomed. Opt. Express 3(12), 3223–3230 (2012).

8. J. Cassidy, A. Nouri, V. Betz, et al., “High-performance, robustly verified monte carlo simulation with fullmonte,” Journal of Biomedical Optics 23(8), 085001 (2018).

9. R. Yao, X. Intes, and Q. Fang, “Generalized mesh-based Monte Carlo for wide-field illumination and detection via mesh retessellation,” Biomed. Opt. Express 7(1), 171–184 (2016).

10. S. Yan, A. P. Tran, and Q. Fang, “Dual-grid mesh-based monte carlo algorithm for efficient photon transport simulations in complex three-dimensional media,” Journal of Biomedical Optics 24(2), 020503 (2019).

11. A. A. Leino, A. Pulkkinen, and T. Tarvainen, “ValoMC: a Monte Carlo software and MATLAB toolbox for simulating light transport in biological tissue,” OSA Continuum 2, 957–972 (2019).

12. S. Powell and T. S. Leung, “Highly parallel monte-carlo simulations of the acousto-optic effect in heterogeneous turbid media,” Journal of Biomedical Optics 17(4), 045002 (2012).

13. C. J. Zoller, A. Hohmann, F. Forschum, et al., “Parallelized monte carlo software to efficiently simulate the light propagation in arbitrarily shaped objects and aligned scattering media,” Journal of Biomedical Optics 23(6), 1 – 12 – 12 (2018).

14. L. Yu, F. Nina-Paravecino, D. Kaeli, et al., “Scalable and massively parallel Monte Carlo photon transport simulations for heterogeneous computing platforms,” J. Biomed. Opt. 23(1), 010504 (2018).

15. NVIDIA Corp., “Optimization – OpenCL Best Practices Guide,” (2011).

16. Advanced Micro Devices, “AMD Accelerated Parallel Processing, OpenCL Optimization Guide,” (2014).

17. R. Yao, X. Intes, and Q. Fang, “Direct approach to compute jacobians for diffuse optical tomography using perturbation monte carlo-based photon ‘replay’,” Biomed. Opt. Express 9, 4588–4603 (2018).

18. D. Badouel, “Graphics gems,” ch. An Efficient Ray-polygon Intersection, 390–393, Academic Press Professional, Inc., San Diego, CA, USA (1990).

19. M. Shevtsov, A. Soupikov, and A. Kapustin, “Ray-Triangle intersection algorithm for modern CPU architectures,” Proceedings of GraphiCon 11, 33–39 (2007).

20. K. Gupta, J. A. Stuart, and J. D. Owens, “A study of Persistent Threads style GPU programming for GPGPU workloads,” in 2012 Innovative Parallel Computing (InPar), 1–14 (2012).

21. Intel Corp., “The Compute Architecture of Intel Processor Graphics Gen9,” (2015).

22. S. Jacques, “mcxyz software.” https://omlc.org/software/mc/mcxyz/.

23. G. Marsaglia, “Xorshift RNGs,” Journal of Statistical Software, Articles 8(14), 1–6 (2003).

